# VarDCL: A Multimodal PLM-Enhanced Framework for Missense Variant Effect Prediction via Self-distilled Contrastive Learning

**DOI:** 10.64898/2026.03.13.711612

**Authors:** Huiling Zhang, Ganzhang Zheng, Ziqi Xu, Haowen Zhao, Shaozhen Cai, Yuxiang Huang, Zihan Zhou, Yanjie Wei

## Abstract

Missense variants are a common type of genetic mutation that can alter the structure and function of proteins, thereby affecting the normal physiological processes of organisms. Accurately distinguishing damaging missense variants from benign ones is of great significance for clinical genetic diagnosis, treatment strategy development, and protein engineering. Here, we propose the VarDCL method, which ingeniously integrates multimodal protein language model embeddings and self-distilled contrastive learning to identify subtle sequence and structural differences before and after protein mutations, thereby accurately predicting pathogenic missense variants. First, leveraging sequence and structural information before and after mutations, VarDCL generates sequence-structural multimodal features via different language models. It incorporates both global and local perspectives of feature embeddings to provide the model with dynamic, multimodal, and multi-view input data. Additionally, a Self-distilled Contrastive Learning (SDCL) module was proposed to enable more effective information integration and feature learning, enhancing the model’s ability to detect sequence and structural changes induced by mutations. Within this module, the multi-level contrastive learning framework excels at capturing information differences before and after mutations within the same modality; meanwhile, the feature self-distillation mechanism effectively utilizes high-level fused features to guide the learning of low-level differential features, facilitating information interaction across different modalities. The VarDCL framework not only ensures the model’s capacity to learn dynamic changes pre- and post-mutation but also significantly improves cross-modal information interaction between sequence and structure, thereby remarkably boosting the model’s performance in distinguishing pathogenic mutations from benign ones. To validate the effectiveness of VarDCL, extensive experiments were conducted. The ablation study demonstrates that all key components of VarDCL contribute significantly. On an independent test set containing 18,731 clinical variants, VarDCL achieved an AUC of 0.917, an AUPR of 0.876, an MCC of 0.690, and an F1-score of 0.789, outperforming 21 state-of-the-art existing methods. Benchmark analysis shows that VarDCL can be utilized as an accurate and potent tool for predicting missense variant effects.

## 1 Introduction

A missense mutation refers to a genetic alteration that changes a single amino acid within a protein’s sequence, and it ranks among the most common types of genetic variations. Such variants can exert a significant influence on protein functions, resulting in a broad spectrum of phenotypic consequences—ranging from harmless to highly pathogenic[1]. In precision medicine, the identification of potentially deleterious variants constitutes a critical task. Although experimental techniques like deep mutational scanning[2] enable high-throughput analysis of variant impacts, they are constrained by drawbacks including long processing times, high expenses, and limited applicability. As a result, the specific characteristics of most variants remain uncharacterized to date.

Against this background, computational prediction models have gradually evolved into indispensable tools. Most of these variant effect prediction (VEP) methods perform analyses by leveraging information such as sequence conservation, population frequency, and the physicochemical properties of amino acids. Rather than relying on a single source of information, integrating results from well-established VEP models can significantly enhance predictive performance[3, 4].

In recent years, two major technological breakthroughs have laid a solid foundation for improving the accuracy of VEP. On one hand, advances in protein structure prediction technologies—with AlphaFold[5, 6] as a representative example—have made it feasible to generate high-precision 3D structure predictions for nearly the entire human proteome. This allows structural features closely linked to how variants affect function—such as relative solvent accessibility and changes in protein stability—to be incorporated into analytical processes. On the other hand, the advent of protein language models (PLMs) has provided robust tools for extracting biological information. Trained on billions of protein sequences or structures, PLMs can capture latent patterns associated with protein structure, function, and evolution, thereby producing rich embedding features to support subsequent analytical work. Driven by these two advances, several cutting-edge VEP methods[7-9] utilize AlphaFold-predicted protein structures to extract relevant physicochemical properties for predicting mutation effects. Meanwhile, Brandes et al.[10] and TransEFVP[11] harness sequence-based PLMs to capture the impacts of mutations. PreMode[12] utilizes sequence-based PLM features, structure-encoded features, and multiple sequence alignment (MSA) features as inputs to determine mutation pathogenicity. However, these VEP methods suffer from several limitations. structure-based VEP methods usually require manual extraction of various biochemical features and cannot fully capture differences in structural properties. In contrast, PLM-based approaches typically derive embeddings exclusively from sequence information, failing to fully acknowledge the decisive role of protein 3D structure, especially the structure before and after mutation. These limitations have severely impeded the interpretation of the clinical effects of missense mutations.

To resolve these challenges, we present VarDCL, a multimodal framework for missense Variant effect prediction via hierarchical self-distilled contrastive learning. The main characteristics of VarDCL are as follows. First, leveraging sequence and structural information before and after mutations, VarDCL generates sequence-structural multimodal features via different language models. It incorporates both global and local perspectives of feature embeddings to provide the model with dynamic, multimodal, and multi-view input data. Additionally, a Self-distilled Contrastive Learning (SDCL) module was proposed to enable more effective information integration and feature learning, enhancing the model’s ability to detect sequence and structural changes induced by mutations. Within this module, the multi-level contrastive learning framework excels at capturing information differences before and after mutations within the same modality; meanwhile, the feature self-distillation mechanism effectively utilizes high-level fused features to guide the learning of low-level differential features, facilitating information interaction across different modalities. The VarDCL framework not only ensures the model’s capacity to learn dynamic changes pre- and post-mutation but also significantly improves cross-modal information interaction between sequence and structure, thereby remarkably boosting the model’s performance in distinguishing pathogenic mutations from benign ones. Evaluated on 18,731 clinical variants, VarDCL achieved state-of-the-art performance with AUC 0.917, AUPR 0.876 and MCC 0.690, surpassing 21 existing VEP methods.

## 2 Materials and Methods

### 2.1 Training and Testing Datasets

We compiled a comprehensive dataset from UniProt, covering 20,516 human protein-coding genes, including 10,538 genes annotated with clinical associations in ClinVar. Pathogenic and likely pathogenic variants were labeled as positive, while benign and likely benign variants were labeled as negative. The dataset includes 89,834 expert-reviewed mutations, consisting of 36,890 positive and 52,944 negative cases. We used February 1, 2021, as a cutoff date; mutations reviewed before this date formed the training set with 71,103 labels, and those reviewed afterward formed the independent test set with 18,731 labels.

### 2.2 Overall Framework of VarDCL

VarDCL is an integrated multimodal framework that accurately predicts the pathogenicity of missense mutations by leveraging the synergistic effects of protein sequence and structural information. The overall architecture of VarDCL is depicted in Fig. 1, which encompasses several key modules: the Initialization Module for Protein Language Model Embeddings, SDCL, and the Classifier Module. These modules work in concert to extract, refine, and classify features from both wild-type (WT) and mutant (MUT) protein sequences and structures, thereby enabling the model to identify pathogenic and benign mutations.

**Fig. 1.**
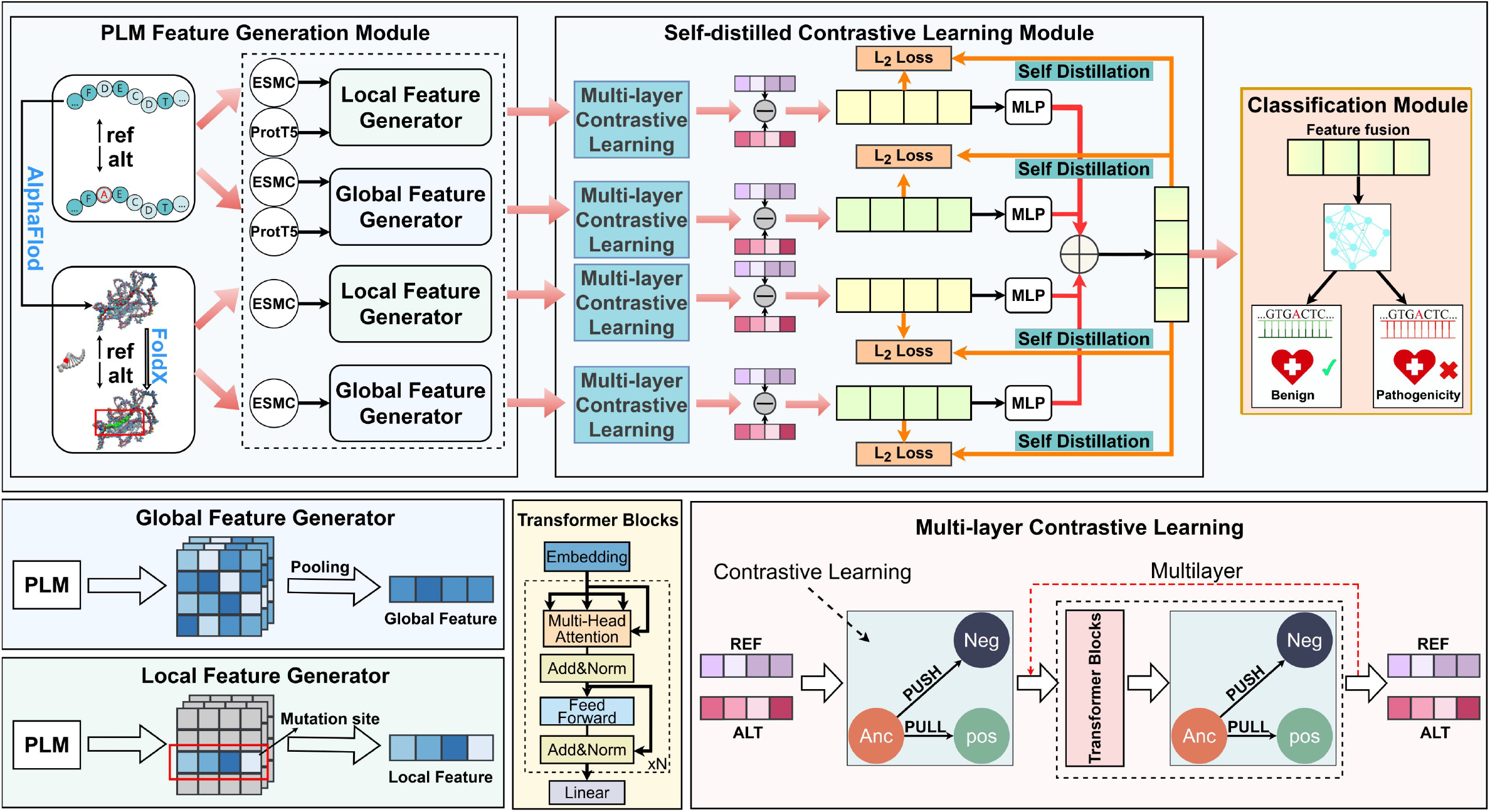
Architecture of VarDCL

### 2.3 Initialization Module for PLM Embeddings in Model Input

The primary task of VarDCL is to generate robust multimodal embeddings for the sequences and structures of wild-WT and MUT proteins. This process integrates various advanced protein language models based on sequences and structures, extracting eight types of embeddings, including global and local embeddings of sequences and structures before and after mutations. Among them, local features refer to the embedding vectors of the mutated residues, while global features are obtained by applying average pooling to the embedding vectors of all residues (for more details, please refer to Appendix A1).

Among numerous language models, we compared the performance of different model embeddings and ultimately selected the two most advanced models: ESMC and ProtT5 (for more details, please refer to Appendix A2). ESMC encodes sequences and structures into 1152-dimensional embeddings, while ProtT5 enriches the contextual information of sequence representations through 1024-dimensional embeddings. The selection of ESMC and ProtT5 is due to their outstanding performance in capturing sequence and structural information, as well as their significant complementarity (for more details, please refer to Appendix A3). These embeddings provide the foundation for downstream prediction modules.

### 2.4 Self-distilled Contrastive Learning (SDCL)

To achieve more effective information integration and feature learning, and to enhance the model’s ability to recognize changes induced by mutations, we introduce a SDCL. SDCL integrates multi-layer contrastive learning, and semantic self-distillation, enabling the model to simultaneously enforce WT and MUT consistency and strengthen sensitivity to subtle, mutation-specific signals.

#### Multi-Layer Contrastive Learning (MLCL)

We propose a multi-layer contrastive learning framework that incrementally extracts and aligns features across different levels within the same modality through multiple layers of contrastive learning. This approach enables a more comprehensive capture of the differences induced by mutations and a deeper understanding of the complex effects of mutations, thereby enhancing the overall performance of the model. The goal of each layer of contrastive learning is to align the representations of WT (wild type) and MUT (mutant) features while maximizing their separation from other samples, thus strengthening the model’s discriminative power. This framework is applicable not only to low-level features but also to high-level semantic features, ensuring effective capture of mutation effects at different levels.

In each layer *n*, the feature representations of the WT and MUT are denoted as 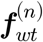 and 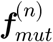, respectively.

These features are extracted from the previous layer’s features using a feature extraction module (such as an improved transformer architecture with multi-head attention and positional encoding (for more details, please refer to Appendix A4), thereby capturing long-range dependencies and deep contextual associations. This process forms a unified semantic foundation that supports deeper contrastive learning and self-distillation.

For each layer *n* and any modality *m* (such as local sequence, local structure, global sequence, global structure), the feature vectors for contrastive learning are mapped to the contrastive space via the projection head ***g*** (⋅):

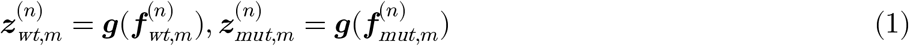

The contrastive loss for modality *m* in the layer*n* is defined as:

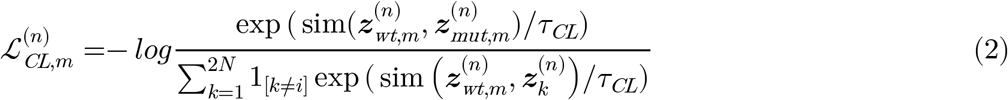

here, sim 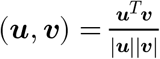 denotes cosine similarity. *𝒯*_*CL*_ is the contrastive learning temperature hyperparameter. 2Ν represents the total number of samples in the sample set, and *K* is used to iterate through all the samples. The 1_*k*[≠ *ⅈ]*_ is an indicator function that takes the value of 1 when *k ≠ ⅈ*, and 0 otherwise. It is used here to ensure that the calculation of the denominator does not include the similarity with itself.

*𝒯*_*CL*_ affects the computation of similarities between samples. A lower temperature amplifies differences, making the model more sensitive to similarities and enhancing its ability to identify mutation features; a higher temperature smooths the calculation of similarities, making the model’s assessment of similarities more nuanced, which aids in learning robust feature representations. For the determination of this parameter value, please refer to 3.3.

The total contrastive loss for each layer is the sum of the contrastive losses across all modalities:

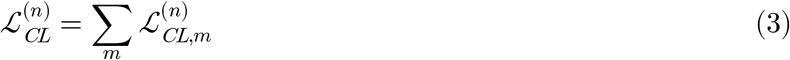

The final contrastive loss of the model is the sum of the contrastive losses from all *Ν* layers:

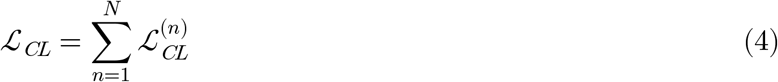

#### Self-Distilled (SD)

Explicitly calculating the differences between WT and MUT features within the same modality is crucial for capturing the effects of mutations. However, relying solely on the differential features within a single modality, without interaction between modalities, may not be sufficient to fully capture the complex effects of mutations. Therefore, we propose an SD mechanism. SD, a special teacher-student model, transfers its knowledge from high-level to low-level blocks[13]. Specifically, we compared several well-known SD methods and ultimately chose the SD method based on L2 norm loss as the SD mechanism for our model (for more details, please refer to Appendix A5). We integrate fusion features (the integration of differential features from each modality) with differential features from different modalities, using the information from high-level fusion features to guide the learning of low-level differential features across modalities. This promotes information interaction between modalities, enabling the model to more sensitively capture subtle structural and sequence variations. This increased sensitivity allows the model to more accurately identify and distinguish between pathogenic and benign mutations, thereby significantly enhancing the performance of the model.

The computation of soft labels plays a key role in this process. Compared to traditional hard labels, which represent a single categorical assignment, soft labels provide a more nuanced way to represent class probabilities. By calculating soft labels, we can more accurately capture subtle distinctions within the data and transfer this information across different levels of the model.

The soft label representation for high-level fusion feature ***f***_*h*_ is computed as:

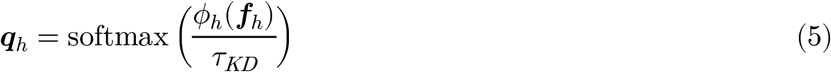

For each low-level differential feature 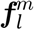, the soft label is computed similarly:

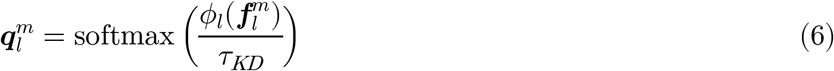

here, *ø* (⋅) is the projection function. The self-distilled temperature parameter *τ*_*KD*_, is used to more sharply adjust the smoothness of the probability distribution, enhancing the effect of knowledge transfer. For the determination of this parameter value, please refer to 3.3.

We utilize mean squared error to minimize the discrepancy between the low-level features and high-level soft labels across various modalities *i* .

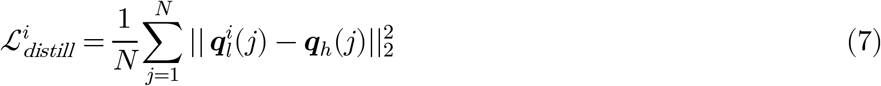

The SD loss is the summation of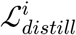:

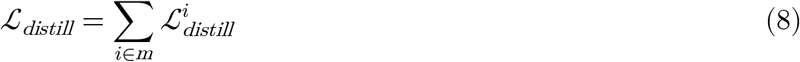

In summary, SDCL not only ensures the ability to learn dynamic changes before and after mutations but also effectively enhances the interaction of information between different modalities, such as sequence and structure, thereby significantly improving the model’s performance in identifying pathogenic and benign mutations.

### 2.5 Classifier Module

To select a robust classifier suitable for high-dimensional features, we conducted experiments on the impact of different classifiers on model performance and ultimately adopted the Kolmogorov–Arnold Network (KAN) framework. This framework replaces the fixed activation operations used in traditional Multilayer Perceptron (MLP) with learnable functional bases, providing a more favorable balance between parameter efficiency and nonlinear modeling capability. Additionally, within this framework, we experimented with the impact of different numbers of KAN-Linear layers on model performance and ultimately determined that the classifier consists of two sequential KAN-Linear layers with output dimensions of 32 and 1. Each layer includes a learnable linear projection, a SiLU nonlinear activation function, and batch normalization to enhance training stability and representational flexibility. To mitigate overfitting, a dropout rate of 0.1 is applied after each of the first two layers. Overall, this lightweight architecture effectively captures the fine-grained nonlinear patterns inherent in biological feature spaces.

### 2.6 Multi-Loss Joint Optimization

The final loss function of our model comprises two components: the Binary Cross-Entropy (BCE) loss for the primary classification task, SDCL. By jointly optimizing these losses, the model learns effective classification representations while simultaneously enhancing the discriminative power of its features. The total loss function is defined as:

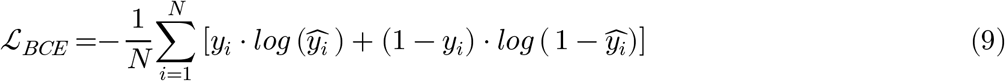

here, *N*is the number of samples, *y*_*i*_ is the true label of sample *i* (0 for benign, 1 for pathogenic), and 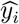 the model’s predicted probability of the variant being pathogenic.

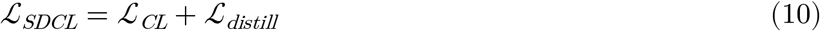

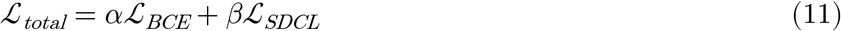

where *α* and *β* are hyperparameters that balance the contribution of the different loss terms.

## 3 Results and Analysis

VarDCL was trained with 10-fold cross-validation. The experiments in this section are conducted on the independent test. To evaluate the performance VarDCL, we employed a suite of key metrics, including the area beneath the receiver operating characteristic curve (AUC-ROC), the area under the precision-recall curve (AUPR), Matthews correlation coefficient (MCC), accuracy, F1 score, precision, recall(sensitivity), and specificity. For implementation, we adopted PyTorch:2.6.0, and the model training and testing were carried out on two Nvidia RTX4090 GPUs equipped with 24 GB of memory.

### 3.1 The Performance of PLMs with Different Modalities and Their Combinations

To select appropriate PLMs for VarDCL to encode sequence and structural inputs, we first evaluated the performance of different sequence-based PLMs (ESM1b[14], ESM1v[15], ESM2[16], ESMC[17], ESM3[17], ProtT5[18]) and structure-based PLMs (ESMC, ESM3). Based on their performance in our missense mutation effect prediction framework, we ultimately selected ProtT5 for generating protein sequence features and ESMC for protein structural feature extraction. For more details, please refer to Fig. A1 in Appendix A2.

Furthermore, we evaluated the effectiveness of different modal features encoded by ProtT5 and ESMC, as well as their combinations, on an independent test set. As shown in Table 1, single-modal features exhibited poor performance—with the lowest performance observed when only ESMC sequence embeddings were used—while significant performance improvements were achieved when combining ESMC sequence embeddings with either ProtT5-encoded sequence embeddings or ESMC-encoded structural embeddings. The multimodal fusion of ESMC and ProtT5 sequence features with ESMC structural features yielded the optimal performance (AUC = 0.917, AUPR = 0.876, MCC = 0.689, F1-score = 0.787), indicating that sequence and structural information are complementary. This multimodal approach integrates features from diverse sources, providing more comprehensive biological insights and significantly enhancing predictive accuracy.

**Table 1.** The impact of different modules in SDCL on VarDCL’s performance

| Method | AUC | AUPR | MCC | Acc | F1 | Pre | Recall | Spe |
| --- | --- | --- | --- | --- | --- | --- | --- | --- |
| Struct (ESMC)+Seq (ProtT5+ESMC) | **0.917** | **0.876** | **0.689** | **0.863** | 0.787 | **0.838** | 0.742 | **0.926** |
| Struct (ESMC)+Seq (ProtT5) | 0.916 | 0.874 | 0.688 | 0.862 | **0.788** | 0.830 | 0.749 | 0.921 |
| Struct (ESMC)+Seq (ESMC) | 0.909 | 0.865 | 0.669 | 0.854 | 0.772 | 0.829 | 0.722 | 0.923 |
| Struct (ESMC) | 0.909 | 0.864 | 0.662 | 0.850 | 0.772 | 0.804 | 0.743 | 0.906 |
| Seq (ProtT5+ESMC) | 0.910 | 0.867 | 0.666 | 0.851 | 0.778 | 0.792 | **0.765** | 0.896 |
| Seq (ProtT5) | 0.905 | 0.858 | 0.658 | 0.848 | 0.770 | 0.797 | 0.744 | 0.902 |
| Seq (ESMC) | 0.897 | 0.845 | 0.641 | 0.842 | 0.756 | 0.799 | 0.717 | 0.907 |

### 3.2 The Impact of Self-distilled Contrastive Learning on VarDCL’s Performance

The SDCL framework of VarDCL consists of two core components, namely the MLCL and SD modules introduced in the Methods section. Among them, MLCL can better capture the information differences within the same modality before and after mutation; the SD module can effectively leverage high-level fused features to guide the learning of low-level differential features, facilitating information interaction between different modalities. To evaluate the impact of different modules in SDCL on VarDCL’s performance, we performed ablation studies on the independent test set (Table 2). Experimental results indicate that removing the MLCL module will reduce the performance of the VarDCL model to a certain extent: the AUC decreased to 0.915 (0.2% drop), and the AUPR decreased to 0.873 (0.3% drop). This result indicates that the MLCL, by progressively extracting and aligning features at different levels, helps the model to more comprehensively capture the complex effects of mutations, enhancing the model’s understanding and discriminative ability regarding mutation effects, thereby improving the overall performance and robustness of the model. Ablation of the distillation module exerts a significant impact on the model: AUC to 0.902 (1.5% decrease), AUPR to 0.855 (2.1% decrease), ACC to 0.835 (2.8% decrease), and MCC to 0.645 (4.4% decrease). This confirms that the SD mechanism makes it possible to dynamically quantify mutations, facilitating the interaction of information about pre- and post-mutation differences between different modalities, therefore enhancing the model’s ability to learn dynamic changes before and after mutations. The SDCL framework, integrated with the MLCL and SD modules, significantly enhances the model’s ability to quantify mutation effects through the effective learning of multimodal information before and after mutations.

**Table 2.** The impact of different modules in SDCL on VarDCL’s performance

| Method | AUC | AUPR | MCC | Acc | F1 | Pre | Recall | Spe |
| --- | --- | --- | --- | --- | --- | --- | --- | --- |
| VarDCL | **0.917** | **0.876** | **0.689** | **0.863** | 0.787 | **0.838** | 0.742 | **0.926** |
| w/o MLCL | 0.915 | 0.873 | 0.687 | 0.861 | **0.790** | 0.815 | 0.766 | 0.910 |
| w/o SD | 0.902 | 0.855 | 0.645 | 0.835 | 0.771 | 0.730 | **0.818** | 0.843 |

Overall, SDCL not only ensures the model’s ability to learn dynamic changes before and after mutations but also effectively enhances the interaction of information between different modalities such as sequence and structure. This significantly improves the model’s identification performance of pathogenic and benign mutations.

### 3.3 Performance of Different Hyperparameters in SDCL

Figure 2 illustrates how hyperparameter settings within SDCL impact model performance on the independent test set. We focused on two key hyperparameters: the contrastive learning temperature (*τ*_CL_) and the self-distillation temperature (*τ*_KD_). The two parameters significantly influence performance by affecting similarity sensitivity in contrastive learning and knowledge transfer efficiency in self-distillation.

**Fig. 2.**
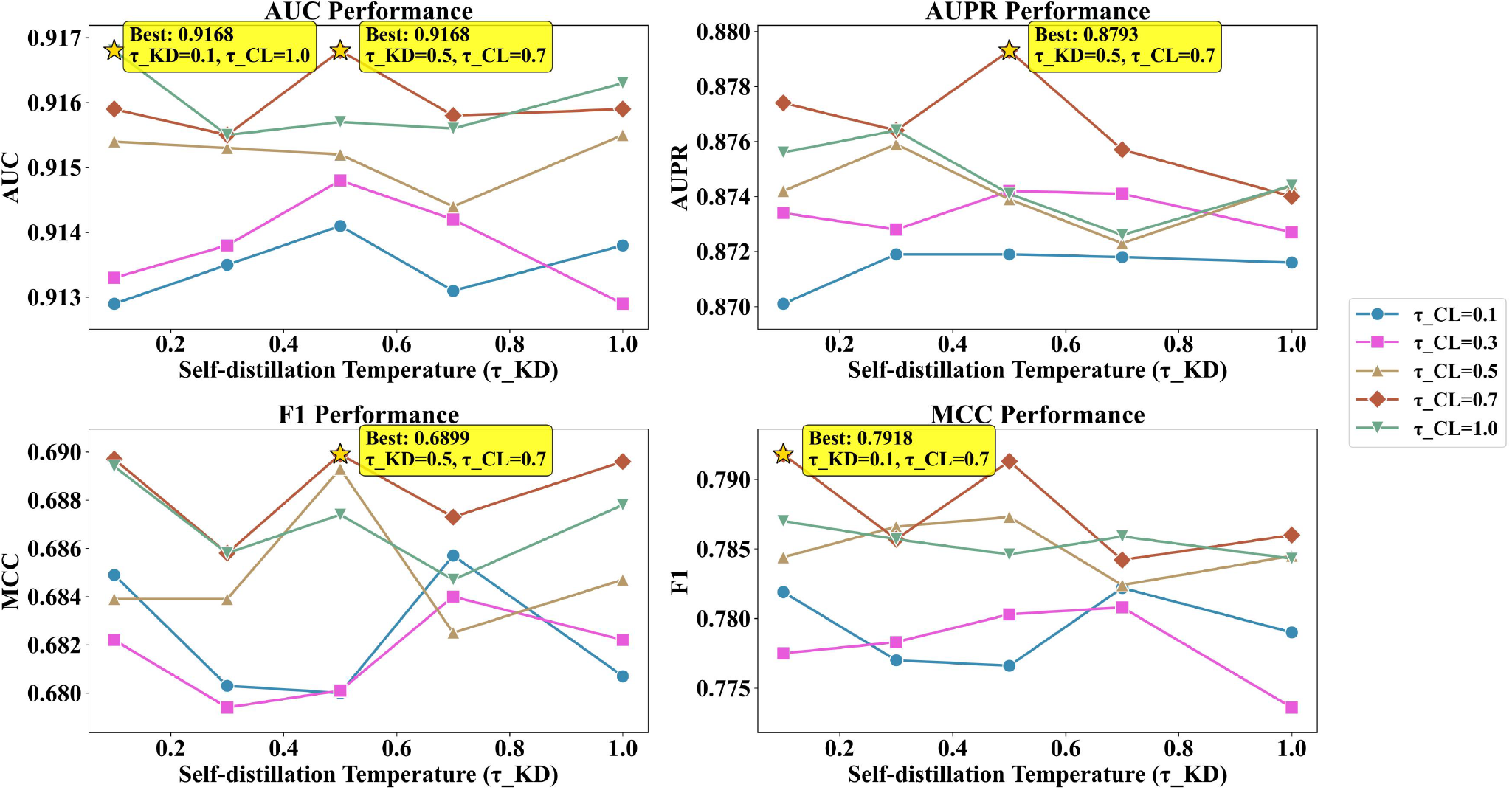
Performance of different hyperparameters in SDCL

*τ*_KD_ controls soft label smoothness in self-distillation. Lower values produce harder labels, potentially increasing noise sensitivity, while higher values improve generalization. *τ*_CL_ affects similarity measurement sensitivity in contrastive loss; lower values enhance mutation characteristic capture, while higher values promote robust feature learning. The results in Fig. 2 show that optimal performance can be achieved for peak AUC (0.917), AUPR (0.879) and F1(0.690) with a moderate *τ*_KD_ (0.5) and a higher *τ*_CL_ (0.7). Aggressive distillation with low temperature degrades all metrics, indicating excessive knowledge transfer can harm representation diversity, while moderate transfer maintains mutation characteristics and enhances discrimination boundaries.

### 3.4 Performance of Different Classifiers

Figure 3 presents a comparison of the performance of various classifiers on the independent test set. Fig. 3(a) compares the performance of multiple predictors, including CatBoost, XGBoost, LightGBM, RandomForest, MLP, and KAN. The results indicate that the KAN model outperforms others across all key metrics, particularly achieving an AUC of 0.876 and an AUPR of 0.863. This indicates that the KAN model exhibits superior performance across diverse scenarios while achieving an excellent balance between precision and recall. Fig. 3(b) further explores the influence of the number of layers in KAN on VarDCL’s performance. The results show that the precision and specificity improve to some extent with an increase in the number of layers, but the two-layer structure performs best across multiple important metrics in AUC (0.917), AUPR (0.879), MCC (0.598), accuracy (0.863), F1 (0.771) and recall (0.729).

**Fig. 3.**
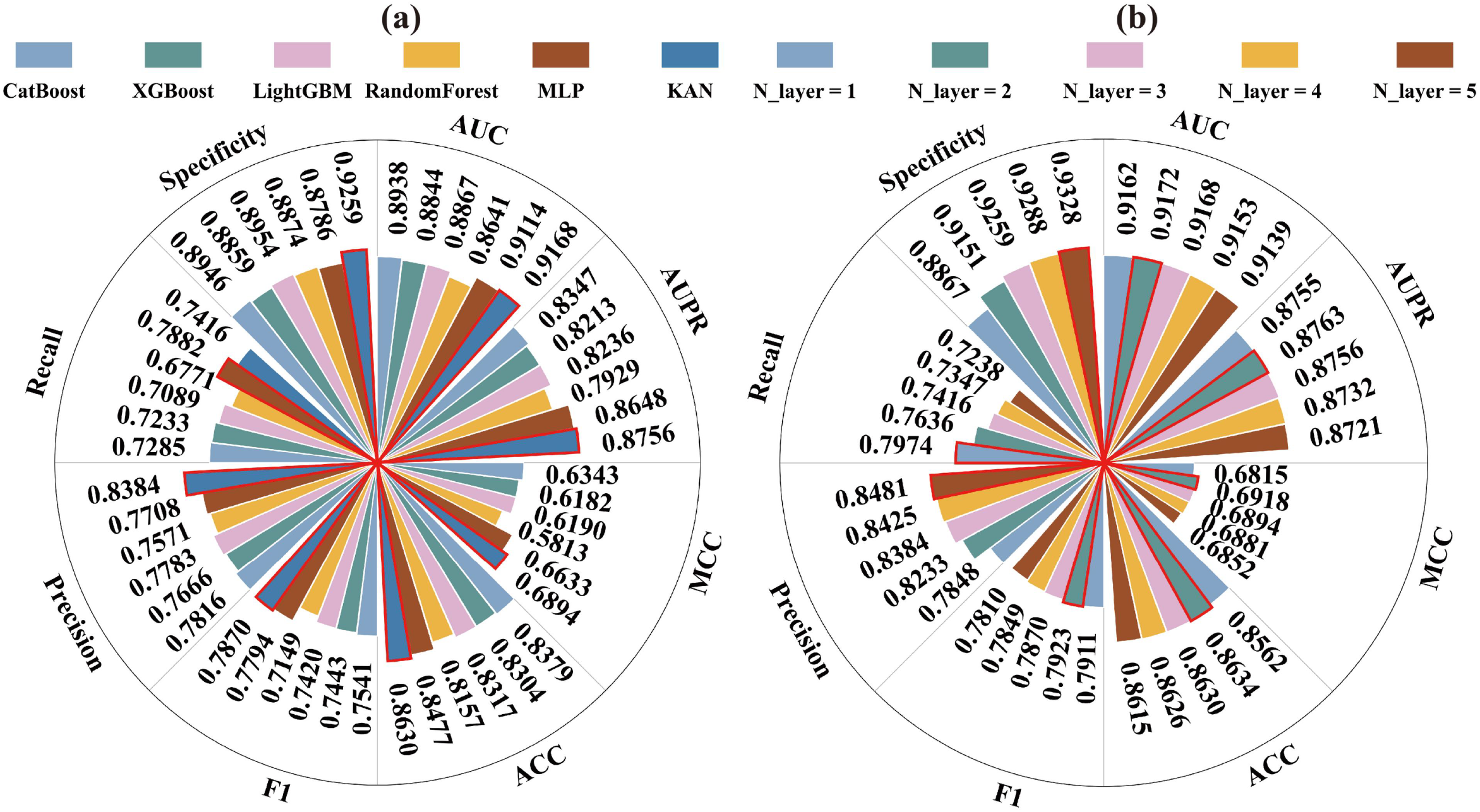
Performance comparison with different predictors

In summary, the results in Fig. 3 indicate that the KAN model not only sets new benchmarks in AUC and AUPR but also leads comprehensively in multiple key performance indicators, further proving its excellent performance and practical value in mutation prediction tasks.

### 3.5 VarDCL Outperforms Existing Methods in Prediction Ability

To evaluate the performance of VarDCL, we compared it with 21 existing variant effect prediction (VEP) methods across a variety of metrics. These methods include PolyPhen2[19], MutationAssessor[20], VEST4[21], DANN[22], REVEL[3], DEOGEN2[23], Eigen[24], FATHMM[25], MetaSVM[26], MetaLR[26], M-CAP[27], LIST-S2[28], PrimateAI[29], MVP[30], gMVP[31], Brandes et al.[10], AlphaMissense[32], AlphaScore[8], CADD[33], SIGMA[9] and TransEFVP[11]. The prediction results of AlphaMissense, gMVP, Brandes et al.[10], SIGMA, and AlphaScore are all derived from the official data provided in their respective papers; the prediction results of TransEFVP were obtained by locally running the source code; and the prediction results of other methods are retrieved from the dbNSFP database [23].

Figure 4 illustrates VarDCL’s superior predictive capabilities. Fig. 4(a) and (b) show that VarDCL achieved an AUC of 0.917 and an AUPR of 0.876, demonstrating its strong ability to distinguish pathogenic and benign mutations under class imbalance. Both ROC and precision-recall curves consistently highlight VarDCL’s superiority, reinforcing its robustness in capturing complex pathogenic signals that unimodal or static encoding strategies might miss. Detailed performance metrics in Fig. 4(c) further confirm VarDCL’s comprehensive excellence. It leads in accuracy (0.863), precision (0.838), and recall (0.742), showcasing balanced performance in correctly identifying both positive and negative cases. With a specificity of 0.926 and an F1 score of 0.787, its predictions are both accurate and effective. Notably, VarDCL’s MCC of 0.689 is the highest among all compared methods, indicating the strongest correlation between its predictions and actual outcomes. These results confirm VarDCL as a leading predictor for variant effect analysis, offering a reliable and compelling solution for practical applications in genomics and precision medicine.

**Fig. 4.**
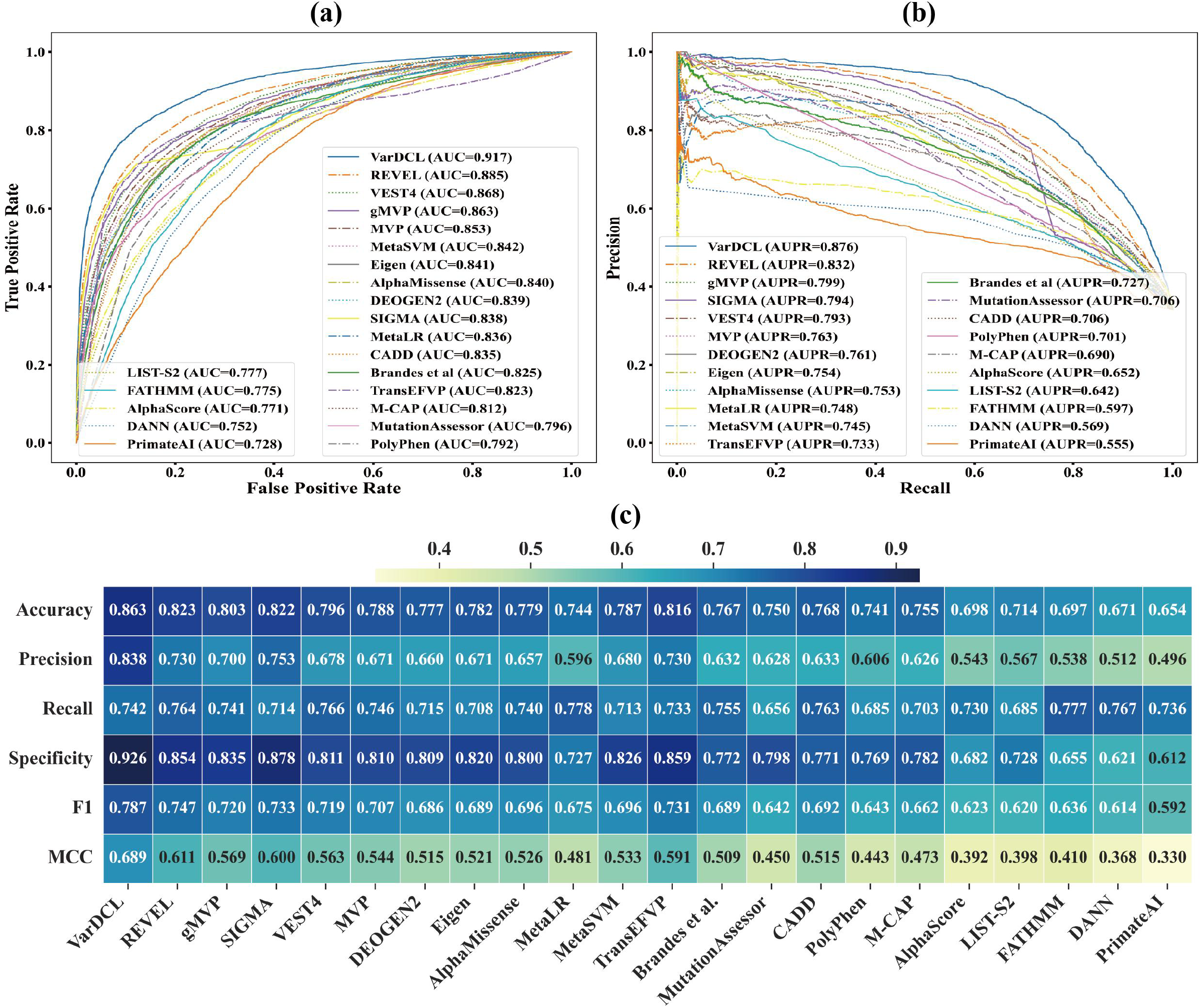
Overall performance comparison of VarDCL with 21 peer methods.

## 4 Conclusions

The core significance of missense mutation effect prediction lies in accurately distinguishing the pathogenicity and benignity of mutations, quantifying their impacts on protein structure and function, thereby providing critical bioinformatics support for genetic disease diagnosis, drug target discovery, and the development of precision medicine strategies. In this study, we propose VarDCL, a method that integrates multimodal protein language model embeddings with self-distilled contrastive learning to capture subtle sequence and structural changes in proteins pre- and post-mutation, enabling accurate prediction of pathogenic missense variants. The SDCL framework of VarDCL comprises two core modules: MLCL (Multi-Level Contrastive Learning) and SD (Self-Distillation). The former accurately captures intra-modal information differences before and after mutations, while the latter guides the learning of low-level differential features through high-level fused features, facilitating cross-modal information interaction. Ablation studies on the independent test set demonstrate that removing either the MLCL module or the SD module leads to a significant decline in all evaluation metrics, confirming that the synergistic effect of the two modules is crucial for enhancing the model’s ability to quantify mutation effects. VarDCL achieved state-of-the-art performance (AUC 0.917, AUPR 0.876, MCC 0.690) on 18,731 clinical variants, outperforming 21 existing methods. Despite these advancements, VarDCL has inherent limitations. Its performance on ultra-rare variants remains suboptimal due to the scarcity of annotated data for such cases, and it relies on the structural accuracy of AlphaFold, which may introduce biases for proteins with complex or poorly predicted 3D structures. Future work will address these gaps by integrating multi-omics constraints (e.g., transcriptomic and epigenomic data) to enrich feature representations, exploring ensemble structure sampling for intrinsically disordered regions to mitigate structural prediction uncertainties, and extending the framework to cross-species generalization to enhance its applicability across diverse organisms. Additionally, we plan to optimize the model for low-resource scenarios, enabling accurate variant effect prediction even with limited annotated data. In summary, by bridging sequence and structural information through synergistic multi-modal learning and self-distilled contrastive mechanisms, VarDCL sets a new benchmark for missense variant effect prediction, offering a reliable and efficient tool to accelerate genetic research, improve clinical decision-making, and facilitate the realization of precision medicine.

## Supplementary Materials

### Appendix A1. Initialization Details of Multimodal Features Initial

#### PLM Embeddings

The wild-type (WT) protein sequence and variant-type (MUT) protein sequence are encoded through the sequence or structure-based PLMs to generate embeddings. All types of embeddings are used for subsequent modules.

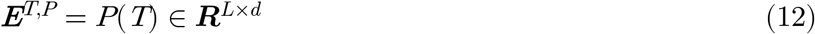

where *T* represents the four modalities of sequence and structure before and after mutation, *P* represents the corresponding PLM. ***E***^*T,P*^ represents embeddings from the PLM. *d* is the dimension of the PLM. *L* is the protein sequence length.

#### Local and Global Feature Extraction

The local feature is the embedding vector of the mutated residue:

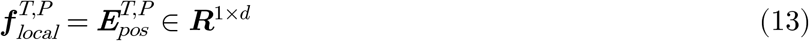

where *pos*(1 *≤pos*≤*L* )is the position index of the mutated residue.

Global Feature is obtained by applying average pooling over the embedding vectors of all residues:

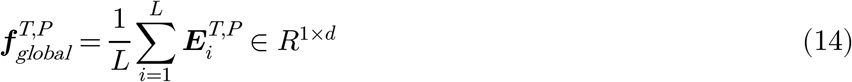

where *i(*1 ≤*i ≤L)* is the position index of the residue.

### Appendix A2. The Selection of Protein Language Models

**Fig. A1.**
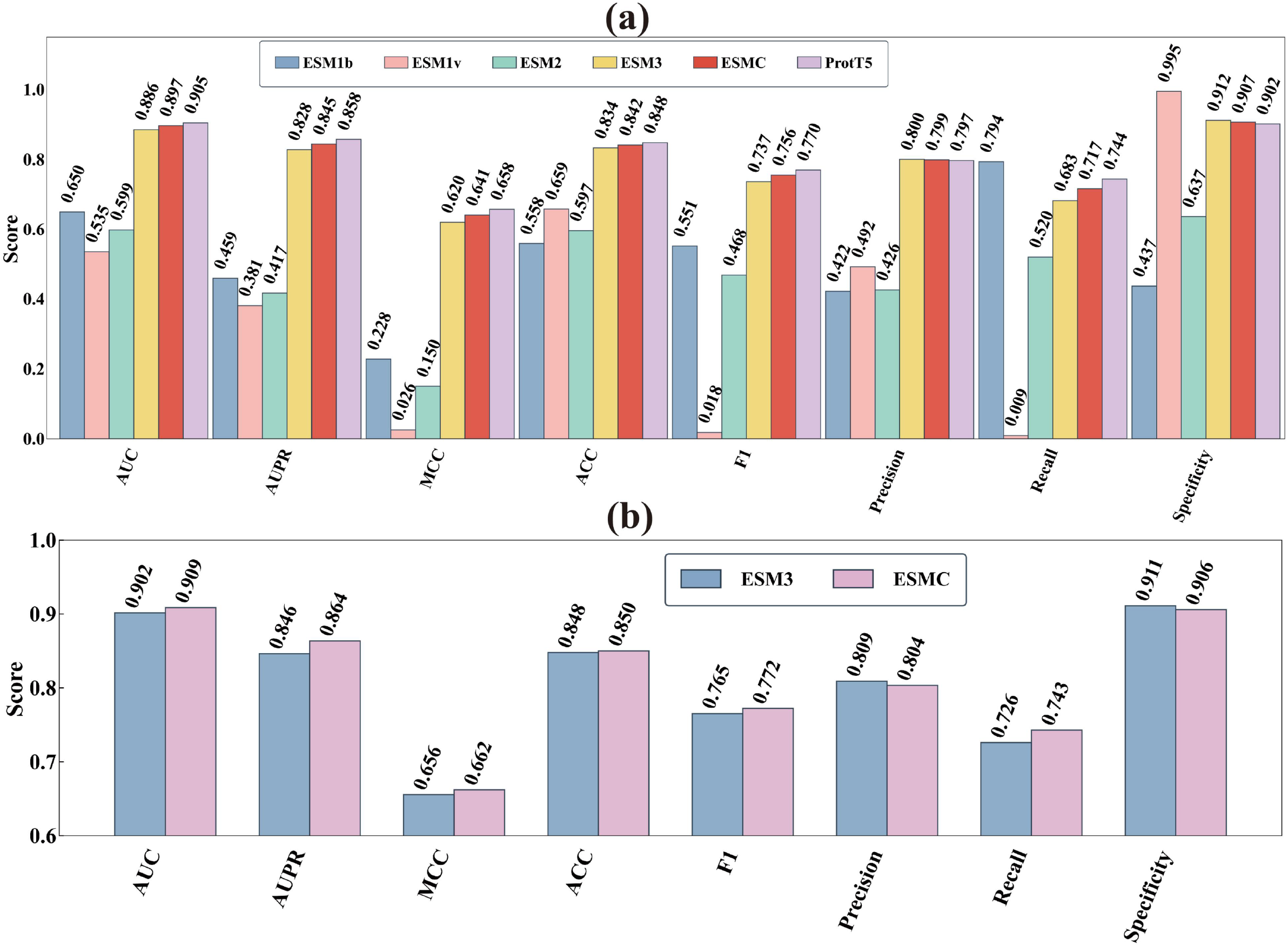
The Impact of Embeddings from Different Protein Language Models on Sequence and Structure Modality for Model Performance

We employed a variety of advanced Protein Language Models (PLMs), including ESM1b, ESM1v, ESM2, ProtT5, ESMC, and ESM3, to generate embeddings for both sequence and structural modalities. These embeddings encompassed global and local features of sequences and structures before and after mutation. For the structural modality, we specifically utilized multimodal protein language models ESMC and ESM3 to generate embeddings. By comparing the impact of embeddings generated by these models on model performance, we found that the ProtT5 and ESMC models demonstrated outstanding performance across both sequence and structural modalities.

Specifically, as shown in Fig. A1(a), ProtT5 excelled in key performance metrics for the sequence modality, including AUC (0.905), AUPR (0.858), MCC (0.657), F1 (0.768), and ACC (0.848), indicating its significant advantage in capturing sequence features. ESMC also performed admirably in the structural modality, as indicated in Fig. A1 (b), outperforming ESM3 in metrics such as AUC (0.909), AUPR (0.864), MCC (0.662), F1 (0.772), and ACC (0.850), demonstrating its efficient capability in capturing structural features.

These results suggest that embeddings generated by ESMC and ProtT5 provide richer and more accurate feature representations for the model, thereby achieving superior performance in predictive tasks. Consequently, we selected ESMC as the baseline model for the structural modality, and chose both ESMC and ProtT5 as the baseline models for the sequence modality to generate the corresponding embeddings, with the aim of enhancing the overall predictive capability of the model.

### Appendix A3. Analysis of Inter-Modality Feature Correlations and Embedding Visualizations

**Fig. A2.**
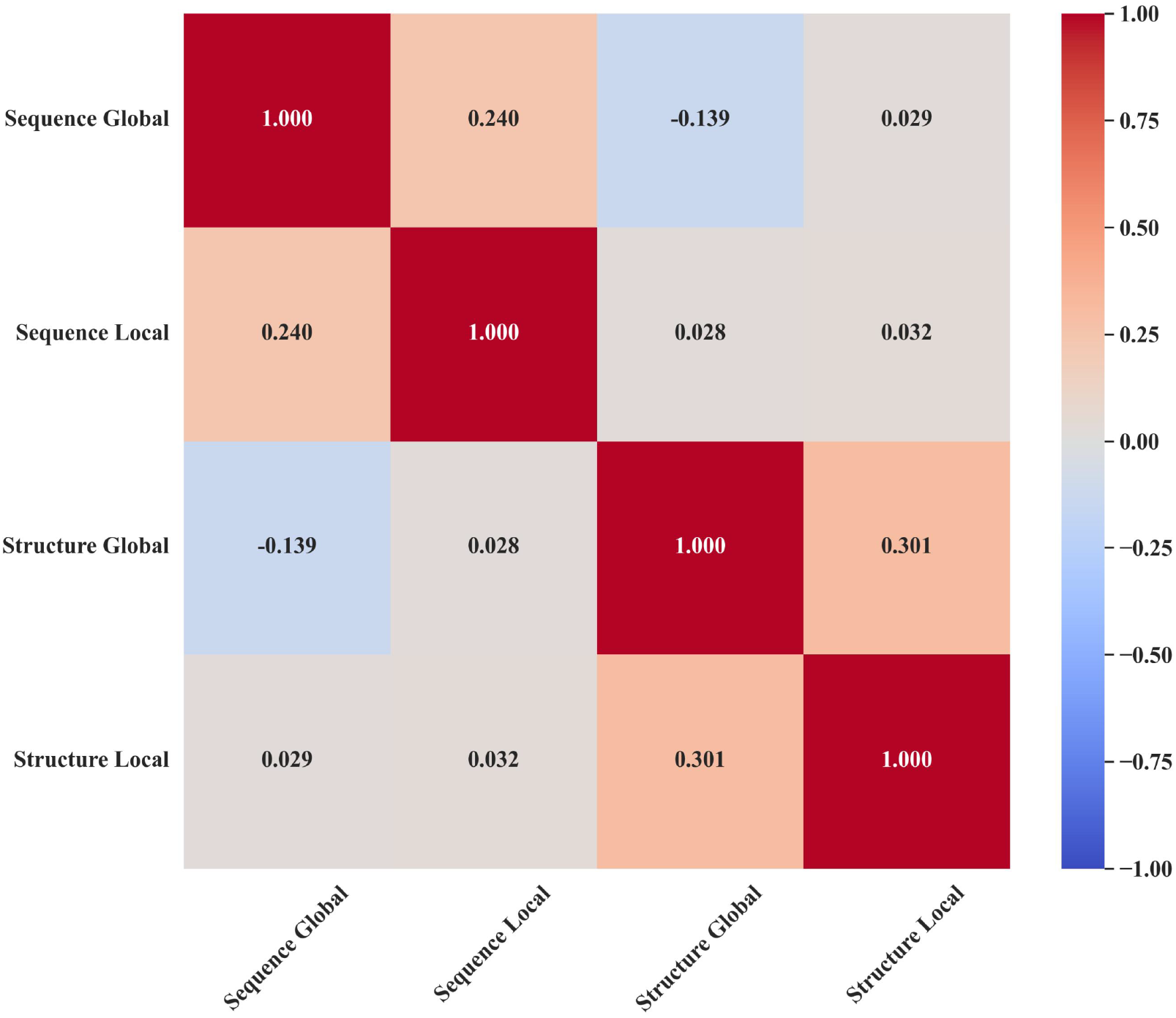
Inter-Modality Feature Correlation Matrix.

Figure A2 illustrates the feature correlations between sequence modalities (Sequence Global and Sequence Local) and structure modalities (Structure Global and Structure Local). Each element in the matrix represents the Pearson correlation coefficient between two modality features, with color intensity indicating the strength and direction of the correlation, where red denotes positive correlation and blue indicates negative correlation. The values on the diagonal are all 1.000, signifying perfect correlation of each modality feature with itself. Off-diagonal values show the correlations between different modality features. For example, the correlation between Structure Global and Structure Local is 0.301, and the correlation between Sequence Global and Sequence Local is 0.240, indicating a moderate positive relationship between them. However, the correlations between sequence modalities (Sequence Global and Sequence Local) and structure modalities (Structure Global and Structure Local) are -0.139 and 0.028± 0.02, respectively, suggesting weak correlations. Nonetheless, the combination of multiple modalities contributes to improved model performance, indicating significant complementary advantages.

**Fig. A3.**
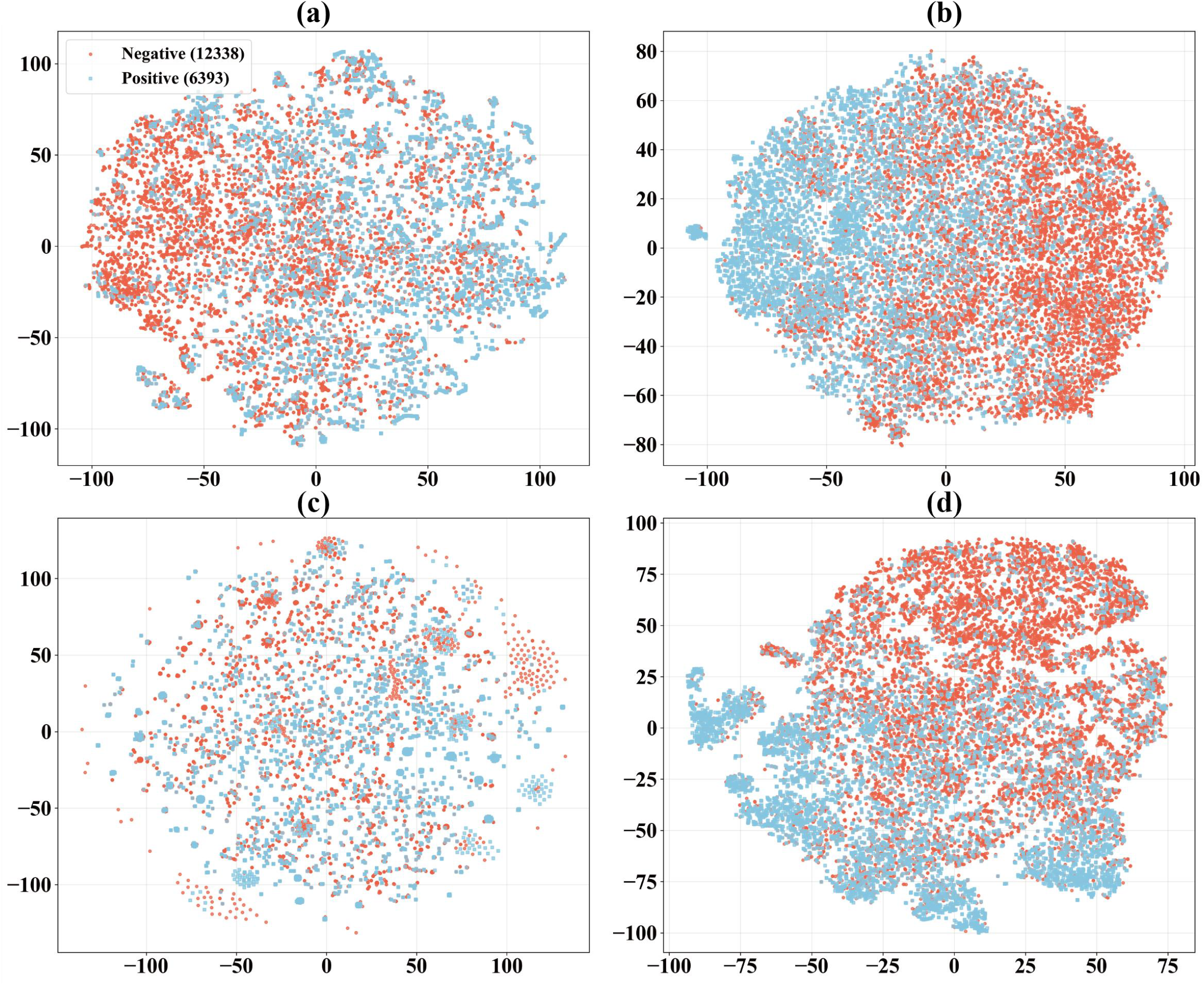
Visualization of the embeddings of Global and Local features: (a) Global embeddings from sequence (b) Local embeddings from sequence (c) Global embeddings from structures (d) Local embeddings from structures.

Figure A3 visualizes the embeddings using t-Distributed Stochastic Neighbor Embedding (t-SNE). t-SNE is a technique for projecting high-dimensional embeddings into two-dimensional space. All t-SNE plots were created with 1000 iterations, a perplexity of 50, and a random state of 42. The visualizations of global sequence, global structure, local sequence, and local structure in Fig. A3 indicate that proteins with similar attributes, whether benign or pathogenic, tend to cluster together. This suggests that the important information about proteins captured by the PLM model, such as sequence, structure, biophysical properties, and biochemical properties, is highly useful for predictive tasks. In other words, this clustering is learned by the pre-trained embeddings, even before seeing labels associated with specific categories. Fig. A3 also shows that there are distributional differences between the embeddings of the sequence and structure modalities, as well as their global and local representations, reflecting their complementary nature in predictive tasks.

### Appendix A4. Performance of Different Hyperparameters in Encoder

**Fig. A4.**
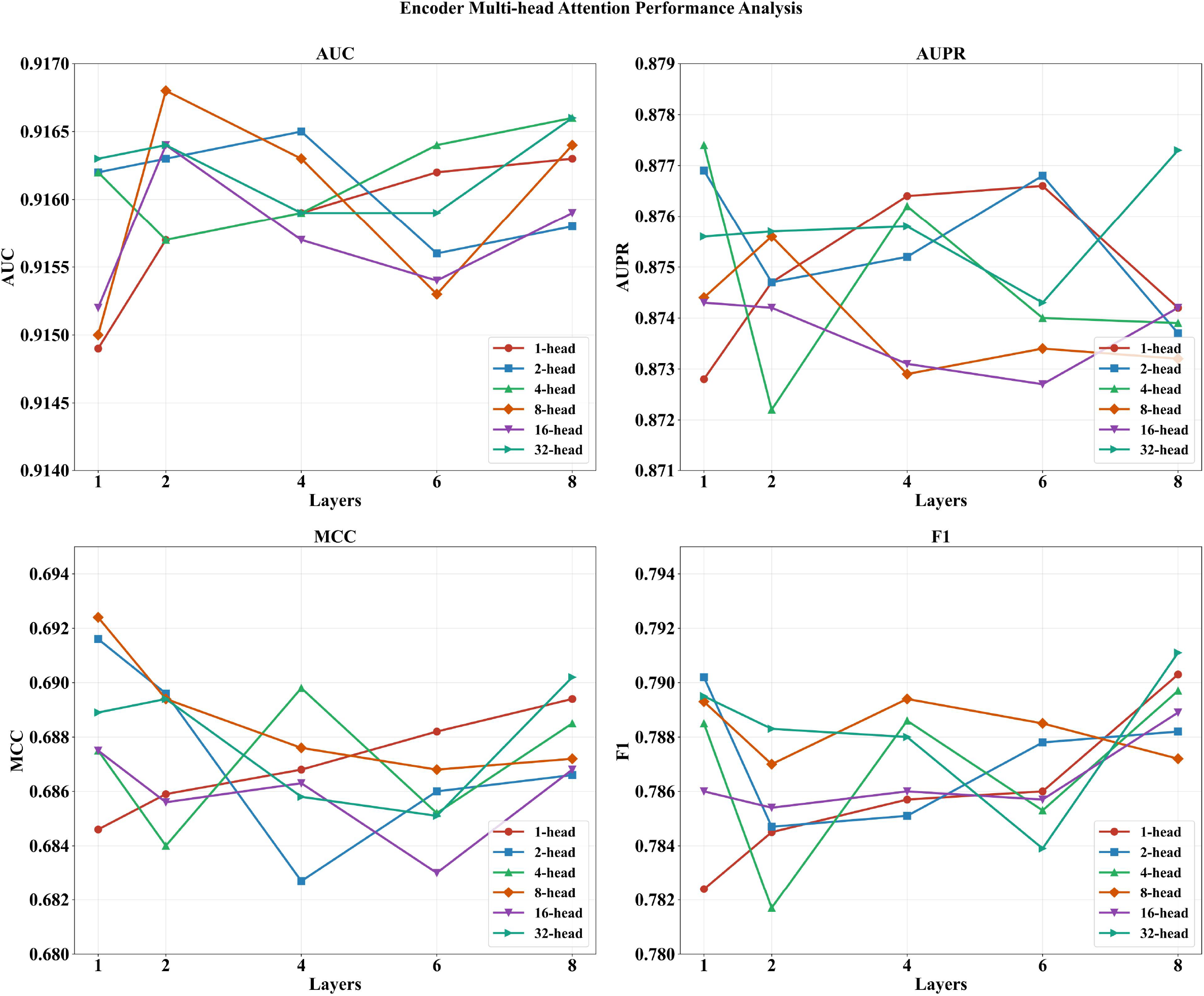
Performance Analysis of Encoder Hyperparameter

In Figure A4, the parameter num_layers denote the number of identical encoder layers stacked, directly influencing the representational depth of the model. The parameter num_heads indicate the number of parallel subspaces in the self-attention mechanism, determining the model’s capacity to capture diverse semantic features. Fig. A4 illustrates the impact of different encoder layer counts and attention head numbers on model performance.

Experimental results demonstrate that the number of heads and layers in the multi-head attention mechanism moderately influences model performance, with MCC and F1-score being more sensitive to architectural changes. The optimal configuration is 8-16 heads combined with 6-8 layers, which demonstrates balanced and stable performance across multiple metrics. However, when the number of attention heads is set to 2, the model achieves optimal performance on the AUC metric, indicating a trade-off between different performance metrics.

### Appendix A5. The Impact of Different Distillation Methods on Model Performance

**Fig. A5.**
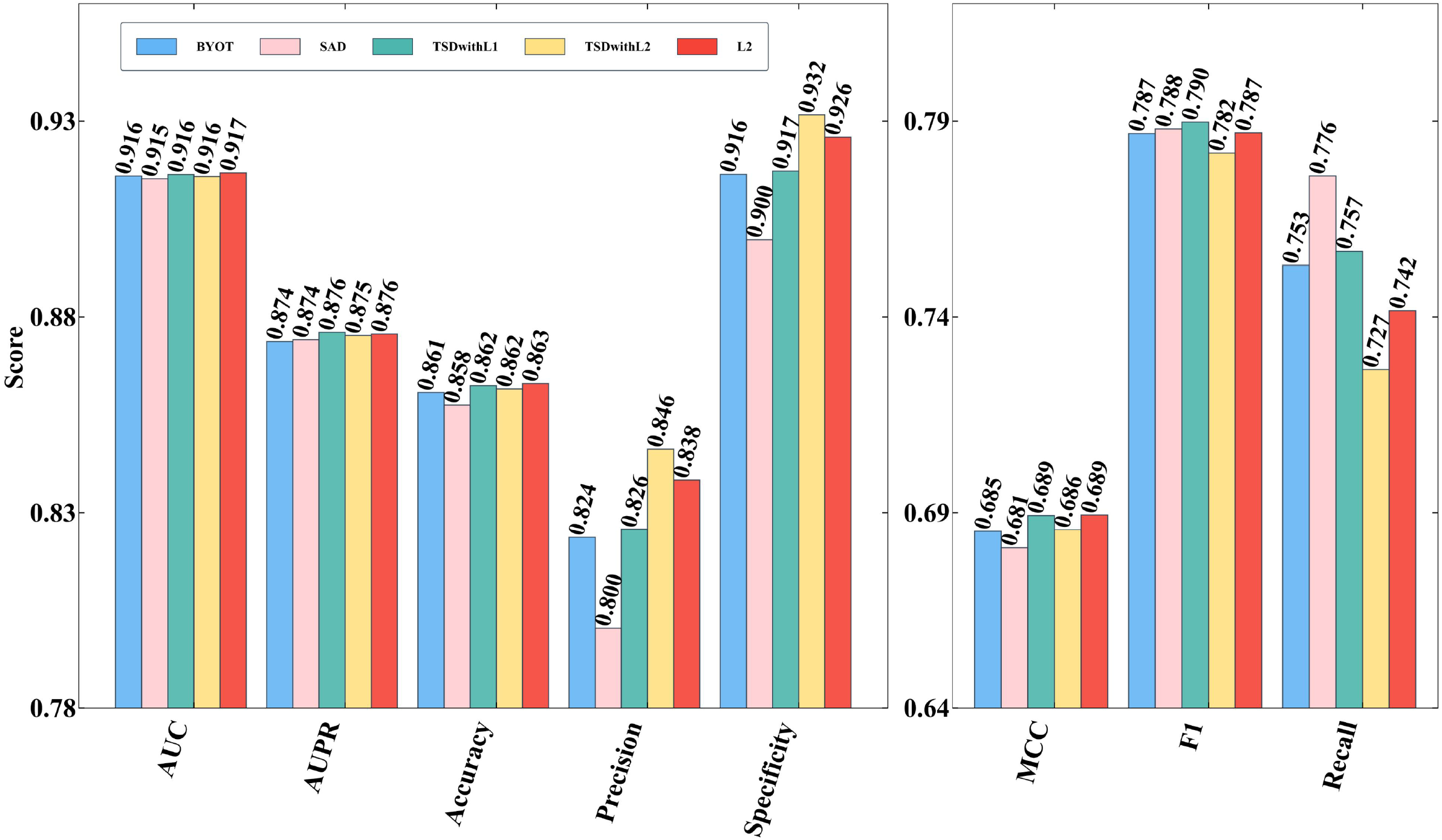
The Impact of Different Distillation Methods on Model Performance

To identify the optimal self-distillation method, we evaluated a selection of five advanced techniques: BYOT, which leverages the Kullback-Leibler (KL) divergence to measure the difference between high-level and low-level feature vectors[13]; SAD, a layer-wise attention-based method that transfers knowledge from higher to lower levels; TSD with KL, a transitive self-distillation approach that uses KL divergence as its loss function[34]; TSD with L2, another transitive self-distillation method that employs the L2 norm as its loss function[34]; and the L2 norm method, which uses the L2 norm to assess the discrepancy between feature vectors at different levels. As shown in Fig. A5, among these, the L2 self-distillation method stood out for its significant positive effect on model performance, particularly in key metrics like AUC, AUPR, MCC, and F1. This method not only enhances the model’s internal knowledge transfer but also strengthens its robustness, leading us to adopt it as our model’s self-distillation approach.

